# Understanding epistatic networks in the B1 β-lactamases through coevolutionary statistical modeling and deep mutational scanning

**DOI:** 10.1101/2023.11.27.568921

**Authors:** JZ Chen, M Bisardi, D Lee, S Cotogno, F Zamponi, M Weigt, N Tokuriki

## Abstract

Over the course of evolution, proteins families undergo sequence diversification via mutation accumulation, with extant homologs often sharing less than 25% sequence identity. The resulting diversity presents a complex view of sequence-structure-function relationships, as epistasis is prevalent, and deleterious mutations in one protein can be tolerated in homologous sequences through networks of intramolecular, compensatory interactions. Understanding these epistatic networks is crucial for understanding and predicting protein function, yet comprehensive analysis of such networks across protein families is limited. In this study, we combine computational and experimental approaches to examine epistatic networks in the class B1 metallo-β-lactamases, a diverse family of antibiotic-degrading enzymes. Using Direct Coupling Analysis, we assess global coevolutionary signatures across the B1 family. We also obtain detailed experimental data from deep mutational scanning on two distant B1 homologs, NDM-1 and VIM-2. There is good agreement between the two approaches, revealing both family-wide and homolog specific patterns that can be associated with 3D structure. However, specific interactions remain complex, and strong epistasis in evolutionarily entrenched residues are not easily compensated for by changes in nearby interactions.

## Introduction

Proteins are a fundamental component of life, and the comprehension of their sequence-structure-function relationship is essential to the understanding and application of a variety of fields, including biology, biochemistry and protein engineering. Over the course of billions of years of evolution, many proteins have emerged to perform a specific function. Subsequently, these proteins further diversified their sequences, forming protein families through accumulation of mutations, *i.e.*, homologous protein sequences spread among diverse organisms. Consequently, homologs often share as little as <25% amino acid sequence identity^1,2^, suggesting evolution has navigated a vast sequence space while maintaining the main protein function. However, the sequence space is often riddled with ‘fitness valleys’^3^. Many recent comprehensive mutant characterization studies, such as deep mutational scanning (DMS), showed that most mutations (∼35-70%) in a single protein are deleterious^4–12^. While variants with these deleterious mutations would have been purged out from the evolutionary process by natural selection, many mutations that are deleterious in a given protein can be observed in other homologous protein sequences, *i.e.*, in the context of many other mutations^13^. These observations suggest that evolution finds mutational paths that go around fitness valleys by exploring networks of intramolecular interactions (*i.e.*, epistatic networks) that can compensate and moderate the deleterious effects of some mutations. In other words, the existence of complex intramolecular networks and coevolution between residues can lead to differential mutational behavior in homologous proteins.

The evolution of natural proteins thus prompts a number of important and unanswered questions. Why can mutations that destroy function in one sequence be fixed in the sequence of homologs? How often are such heterogeneities in mutational effect encountered in a protein family? Can these patterns be modeled and predicted? Knowledge of such epistatic networks within a protein sequence is key to the understanding of the sequence-structure-function relationships, especially in predicting protein sequences that encompass functional proteins. Thanks to advances in both genomics (sequencing)^14^ and biochemical (DMS)^15^ approaches, we can obtain information on the epistatic network from both: as the coevolutionary trend in sequences within an homologous family, or as the experimentally measured mutational behavior in each protein. Epistatic networks can be observed as coevolution of amino acid residues within a protein family^16,17^. Statistical models trained on multiple sequence alignments are able to capture those patterns of coevolution and even reproduce them by generating artificial sequences that respect the statistics of protein families^16,18–22^. While such data-driven computational approaches have demonstrated to be powerful enough to design functional protein sequences that do not exist in nature^23–25^, it is still unclear to what extent and accuracy such models can capture protein epistatic networks in terms of specific interactions. Experimentally, the influence of epistatic networks can be detected as mutational incompatibilities, *i.e.*, mutations that can be tolerated in one genetic background but may be deleterious in another. Such experimental insight provides an in-depth understanding of specific interactions within proteins in terms of biochemical and biophysical mechanisms. However, the experimental study of functional constraints at a family-wide level by comparing residue-level mutational epistasis across homologous sequences remains scarce. This is because only a small number of protein families have been studied^13^, including a few large-scale datasets acquired through DMS^26–31^. Furthermore, most analyses are restricted to higher-level, broader trends between homologs with limited examinations of detailed mechanistic bases for epistasis.

In this study, we combine these two complementary approaches to improve our understanding of epistatic networks. We apply both approaches to the same system, the class B1 metallo-β-lactamases (MBL), which is a family of highly diversified antibiotic degrading enzymes with a long evolutionary history^32^. Using Direct Coupling Analysis (DCA)^16^, we analyze the coevolutionary signatures of the entire B1 family, arriving at a global statistical description of epistatic tendencies based on sequence data. At the same time, we perform DMS on two distantly related members of the B1 family, NDM-1 and VIM-2 (∼30% sequence identity), revealing protein-specific mutational trends and incompatibilities as experimentally measured through functional characterization. By employing both methods in parallel, we have the opportunity to compare the results between them and to learn how they can strengthen and explain each other. We find a general consensus between the two methods on the prevalence and strength of epistasis, and find interesting cases where complementary information provides insight beyond what can be discerned from either method individually. There appears to be trends of mutational behavior in the structure, but the exact intramolecular network appears much more complex upon further testing, as predicted by the DCA model. Overall, the combination of DCA and DMS allows us to greatly enhance our understanding of epistatic networks.

## Results

### Co-evolutionary model of the B1 family reveals sequence-specific mutational heterogeneities

Class B1 metallo-β-lactamases are a highly diversified family of enzymes (as low as 20% identity within the family) that degrade β-lactam antibiotics^1,33,34^, a function for which there is likely a long evolutionary history^32,35^. Given the high sequence divergence, this family is an excellent model for epistatic networks, as different homologs likely formed different sets of intramolecular interactions over the course of their evolution. To create a comprehensive dataset of all B1 sequences, we collected ∼5000 sequences that belong to B1 MBLs, from the MBL domain superfamily (Interpro ID: IPR001279) along with metagenomic data from the Joint Genomics Institute Integrated Microbial Genomes & Microbiomes (JGI IMG/M) database (**see methods**). Then, we inferred a global statistical model using DCA (**methods**) on these B1 MBL homologs, that were aligned in a multiple-sequence alignment (MSA) based on a seed multiple-structure alignment of 14 diverged homologs (**methods**); positions outside the structurally conserved region were excluded. The DCA model captures the coevolutionary relationships (direct couplings) between all pairs of residues in the B1 family, and the residue conservation (fields), that are used to assign a probability *P(sequence)* to any sequence. High probabilities indicate functional MBLs, and low probabilities dysfunctional ones. To score mutations, we take the difference in the model’s “energy” (*E*) between the mutant sequence and the wild-type (WT), as Δ*E* = – log[*P(*mutant)/*P(*WT)]^36^. We can thus predict the effect of any amino acid substitutions from any WT background, with the sequence context taken into account by the coevolutionary couplings.

To probe the mutational behavior across the B1 family, we calculated Δ*E* for all single mutants of 100 diverse homologs, chosen to minimize pairwise sequence identity. To gauge the effects of epistatic networks on each protein position, we measure the mutational tolerance at each position in each homolog as the Shannon Entropy of all mutant probabilities relative to the wild type, also referred to as the context-dependent entropy (CDE)^37,38^. The mutational tolerance (CDE) is the base-2 logarithm of the effective number of tolerated mutations at a position in this specific WT background, such that 0 means only 1 (2^0^) residue is tolerated (the WT, no mutations), and 4.3 means all 20 (2^4.3^) amino acids are equally tolerated. The mutational tolerance at each aligned position in the conserved B1 family is shown for 100 homologs as a heat map in **Fig. 1b**. The figure reveals a number of interesting aspects about the B1 family and its constituent homologs. We can see that there are regions that are highly constrained in terms of mutations, but also others that are highly tolerant. Moreover, homologs closer in the phylogeny show similarity in mutation patterns, and different clades present different patterns.

**Figure 1.**
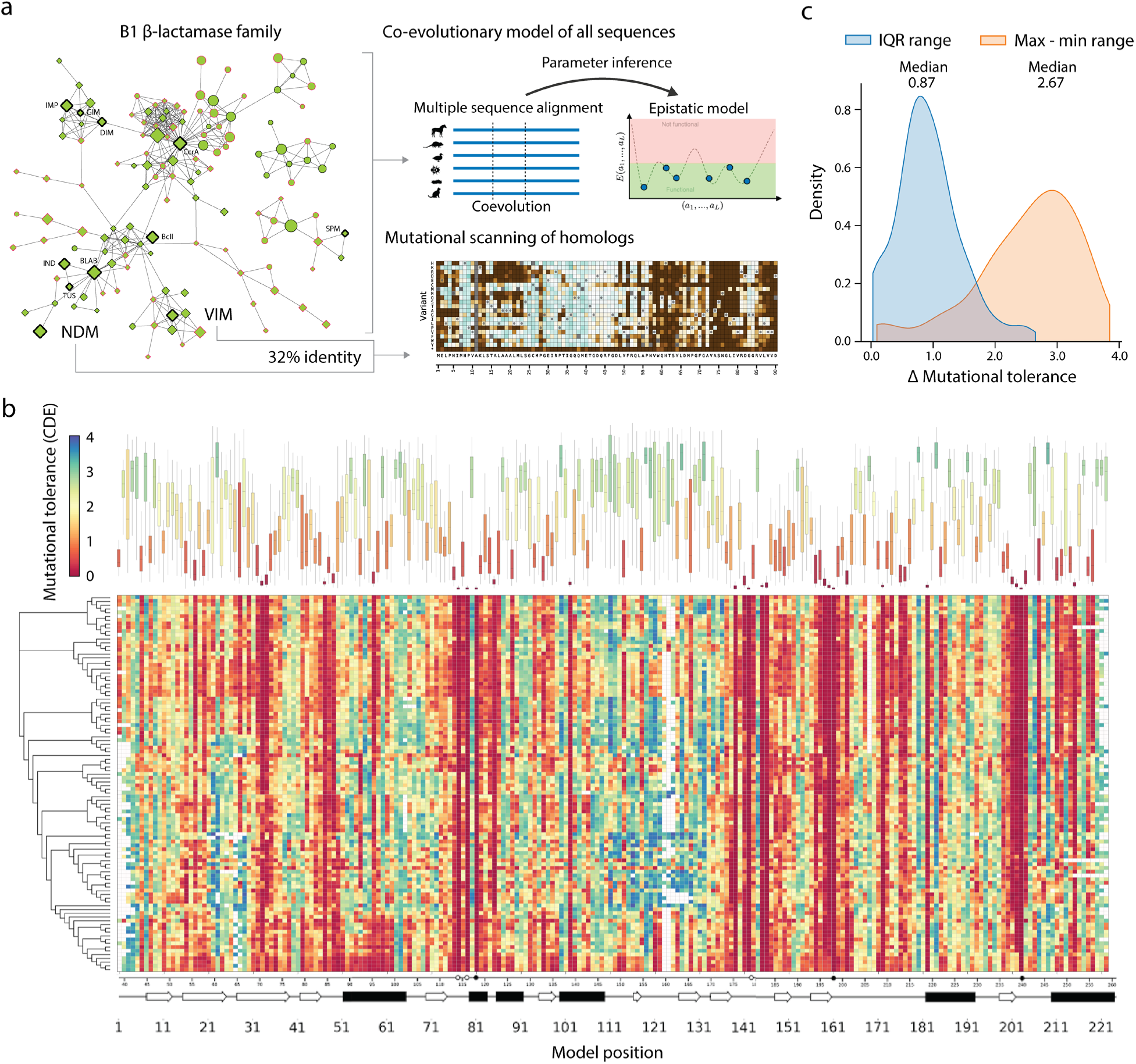
Family-wide residue mutability by direct coupling analysis (DCA). (**a**) Schematic of computational and experimental workflow. The B1 β-lactamase enzyme family is isolated through sequence space exploration via a sequence similarity network. The entire set of sequences are used to generate a co-evolutionary model via DCA. Two highly-diversified homologs within the family are selected for DMS to generate a large mutational dataset. **(b)** Each square in the heat map is colored by the mutational tolerance (measured as context-dependent entropy, CDE) for each position of the 100 aligned homologs. Blank cells represent alignment gaps. For each position the percentiles of the distribution of CDEs are presented as a box plot on top of the heat map: 0-100% as thin lines, 25-75% (IQR) as bars. The bars are colored by the median value, with the same color scale of the heatmap. The secondary structure of the homologs, as well as the active site residues, are depicted under the heat map. The maximum likelihood phylogenetic tree of the 100 homologs is shown on the left of the heat map. **(c)** Distribution of the spread in mutational tolerance across homologs at each position, measured as the distance of the IQR or max – min mutational tolerance.

We highlight substantial heterogeneity in behaviors across the 100 homologs at each position, by quantifying the variability in mutational tolerance at each position as the interquartile range (IQR), (distance from 25-75 percentile) (**Fig. 1b-c**). There is considerable spread of mutational tolerance across homologs in most positions, suggesting that positions that only allow for few mutations in one homolog may show substantial mutability in other homologs. We see a median IQR of 0.87 (2^0.87^=1.83), meaning that half of positions have at least a 1.8-fold difference in effective number of tolerated amino acids between homologs. The IQR per position can be as high as 2.65, or a 6.3 fold difference in effective amino-acid number. Furthermore, the difference in minimum and maximum mutational tolerance across positions has a median of 2.67 and a max spread of 3.85 (**Fig. 1c**), meaning half of the positions in the protein have a 6.3-14.4 fold difference between the most and the least mutationally tolerant sequences. This strong mutational heterogeneity of equivalent positions across homologs is a hallmark of epistasis, and is captured by the coevolutionary couplings in the DCA model.

### Heterogeneity in mutational behavior is supported by DMS data of NDM-1 and VIM-2

To gain experimental observations for differential mutational effects between homologous enzymes, we conducted DMS to obtain all single-mutational effects on two homologs, NDM-1 and VIM-2 (∼30% sequence identity). A complete DMS dataset of all single amino acid mutants of VIM-2 was previously published by us^6,39^. To conduct a comparison, we performed DMS on NDM-1 in an identical manner to VIM-2 (**Fig. 2a**). Briefly, all single amino acid mutants were generated for NDM-1 and placed under selection under three different antibiotics: ampicillin (AMP), cefotaxime (CTX), and meropenem (MEM). The plasmid DNA was isolated after selection and sent for deep sequencing. The fitness conferred by each mutant relative to wtNDM-1 was then characterized as the fitness score in **Eq. (1)**:

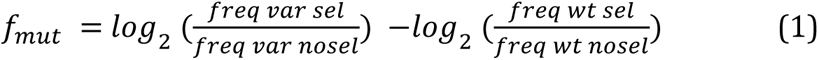

**Figure 2.**
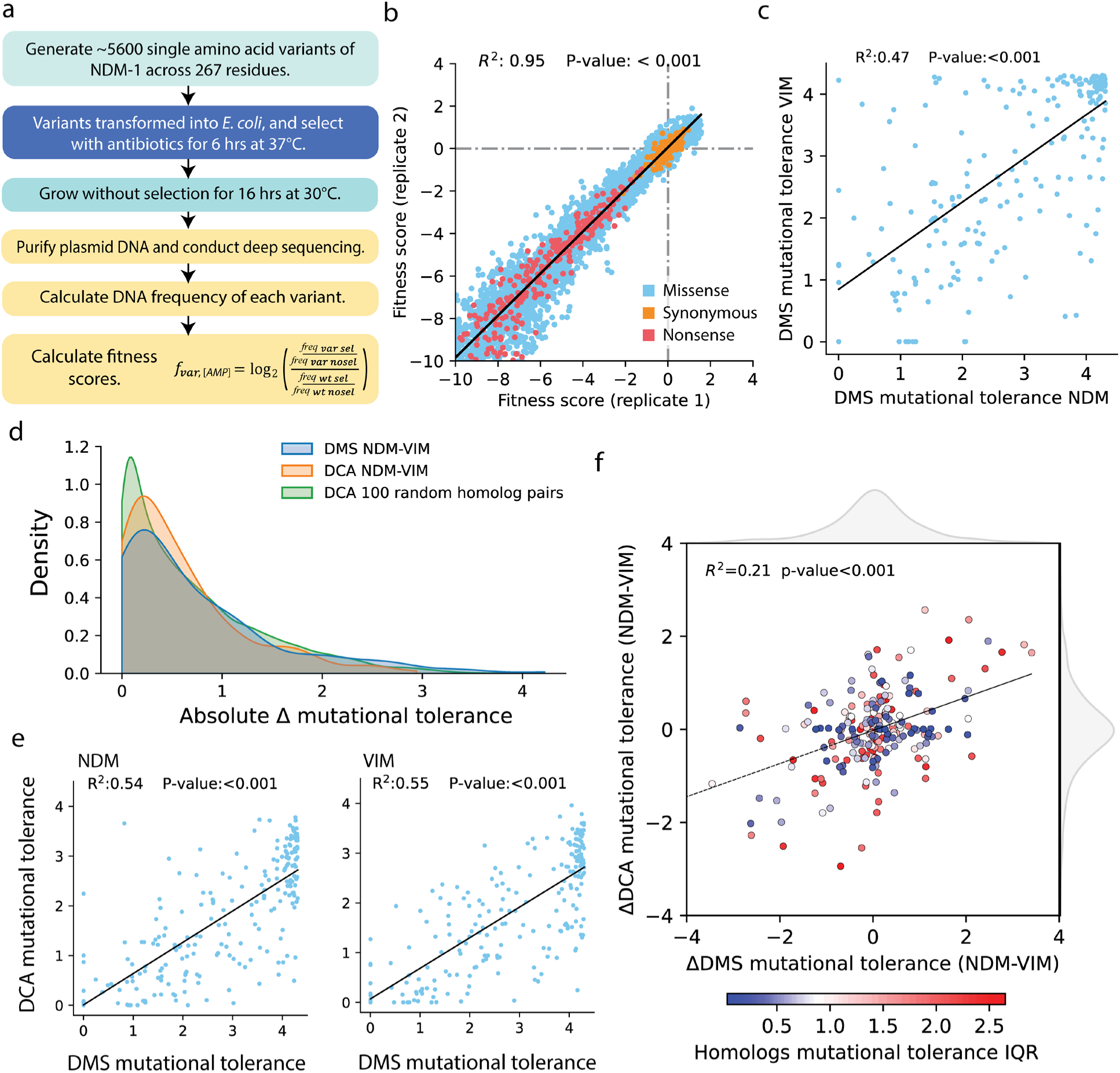
Overview of DMS for NDM-1 and structural similarity to VIM-2. (**a**) Workflow for DMS of NDM-1. (**b)** Correlation between replicates of the NDM-1 library selected at 256µg/mL AMP. The R^2^ and P-value of a linear regression is shown at the top. **(c)** Comparisons of mutational tolerance at each aligned position for the NDM-1 and VIM-2 experiments. The R^2^ and line of best fit for a linear regression are shown. **(d)** Distribution of differences in mutational tolerance in DMS or DCA between NDM-1 and VIM-2 at the same aligned position. The distribution of the difference in DCA mutational tolerance between 100 random pairs of homologs is also plotted. **(e)** Comparison of mutational tolerance between DMS and DCA for NDM-1 and VIM-2. The R^2^ and line of best fit for a linear regression are shown. **(f)** Comparison of the mutational tolerance difference between NDM-1 and VIM-2 at each position between DMS and DCA. The data is colored by the IQR of mutational tolerance across the 100 homologs, with the colors scaled to the distribution (median is the center). The R^2^ and line of best fit for a linear regression are shown.

The experiments were conducted in duplicate on separate days, and replicates typically show good correlation (R^2^ of 0.77-0.95 across conditions) (**Fig. 2b**, **Supp.** Fig. 1). As was the case for VIM-2, the NDM-1 data shows that fitness scores of –4 or lower are below the fitness of nonsense variants. As such, we also limit the fitness scores to a minimum of –4 for analyses, and scores below –4 are set to be –4 instead. To ensure the best possible direct comparison, we use the datasets between NDM-1 and VIM-2 with the most similar selection pressure in AMP, evidenced by the similarity in distribution of fitness effects (**Supp.** Fig. 2).

In a similar fashion as in the DCA analysis, we define the DMS mutational tolerance of each position in NDM-1 and VIM-2 as the Shannon entropy of the mutational probability of each amino acid (**methods**). We find mutational behaviors in the same protein positions to be quite varied between the two homologs, where differences in certain positions can span nearly the full range of possible entropy values, *i.e.*, the same position can accept only the WT amino acid in one homolog (entropy 0) and almost all amino acids in the other homolog (entropy >4) (**Fig. 2c**). We compare the magnitude of entropy differences between VIM-2 and NDM-1 in the DMS data to differences in CDE computed from DCA, and we find strikingly similar distributions (**Fig. 2d**). The median difference between two homologs across all positions is 0.45 in DMS (0.39 in DCA), meaning that half of the positions have more than a 1.37 fold difference in entropies, ranging up to a maximum of 4.22 (18.7 fold). As a baseline for the expected behavior between any pair of homologs, we also computed the CDE difference between 100 random pairs of homologs (**Fig. 2d**). We found a similar distribution, suggesting a common statistical trend of mutational behavior between homologs where certain positions exhibit similar tolerance (such as conserved sites) while others can greatly vary in mutation tolerance.

We further probe whether the DMS and DCA data are in agreement specifically on NDM-1 and VIM-2, by comparing mutational tolerance between DMS and DCA of each position within either NDM-1 or VIM-2 (**Fig. 2e**). There is significant correlation (R^2^∼0.55) between the DMS and DCA values for each of NDM-1 and VIM-2, and this correlation is slightly higher than between the DMS mutational tolerances of NDM-1 vs. VIM-2. This observation shows that DCA captures sequence-specific trends to a good degree. Despite the significant correlation, DCA predicts fewer variable sites (entropy > 3.5) compared to DMS. A possible explanation of this difference may be attributed to the fact that DMS experiments are limited to specific experimental conditions, i.e. a single antibiotic, while DCA likely reflects broader evolutionary constraints. Furthermore, the mutational tolerance differences between VIM-2 and NDM-1 from DCA can also capture the specific differences in DMS (**Fig. 2f**); the correlation is somewhat weakened by the abundance of positions with correctly predicted small differences between NDM-1 and VIM-2. When the variability across homologs (IQR of 100 homologs mutational tolerance distribution) is overlaid on the mutational differences between NDM-1 and VIM-2, we see that positions with the lowest spread in all homologs tend to show lower differences between NDM-1 and VIM-2 in both DMS and DCA, as expected (**Fig. 2f**). Meanwhile, positions with large differences across all homologs are spread throughout. In the case of positions with large differences in either DCA or DMS, the varied behaviors of NDM-1 and VIM-2 match the variation of all homologs. Interestingly, there are also positions that have small differences between NDM-1 and VIM-2, but have high variability across all homologs, further underscoring the fact that mutational behaviors can be quite varied between different homologs. Hence, the combination of methods can reveal and reinforce patterns that would not be obvious from just a single approach.

### Structural basis of mutational tolerance and incompatibilities

We investigate the relationship between mutational heterogeneity and structure, by analyzing the experimental datasets and the model predictions in terms of the protein structure of the B1 family, using the crystal structures of NDM-1 (PDB ID:3spu) and VIM-2 (PDB ID:5yd7) as representatives. For VIM-2 and NDM-1, we quantify the discrepancy in mutability using the absolute difference in mutational tolerance, computed via the DMS experiments and the DCA model, while for the 100 homologs, the IQR of mutational tolerance serves as a DCA-based indicator of variability. It should be noted that points exhibiting either very high or very low average mutability obviously have inherently restricted variability. This is evidenced by the observation that data points in **Fig.3** with a mean or median mutational tolerance lower than 1 or greater than 3 always have low spread.

**Figure 3.**
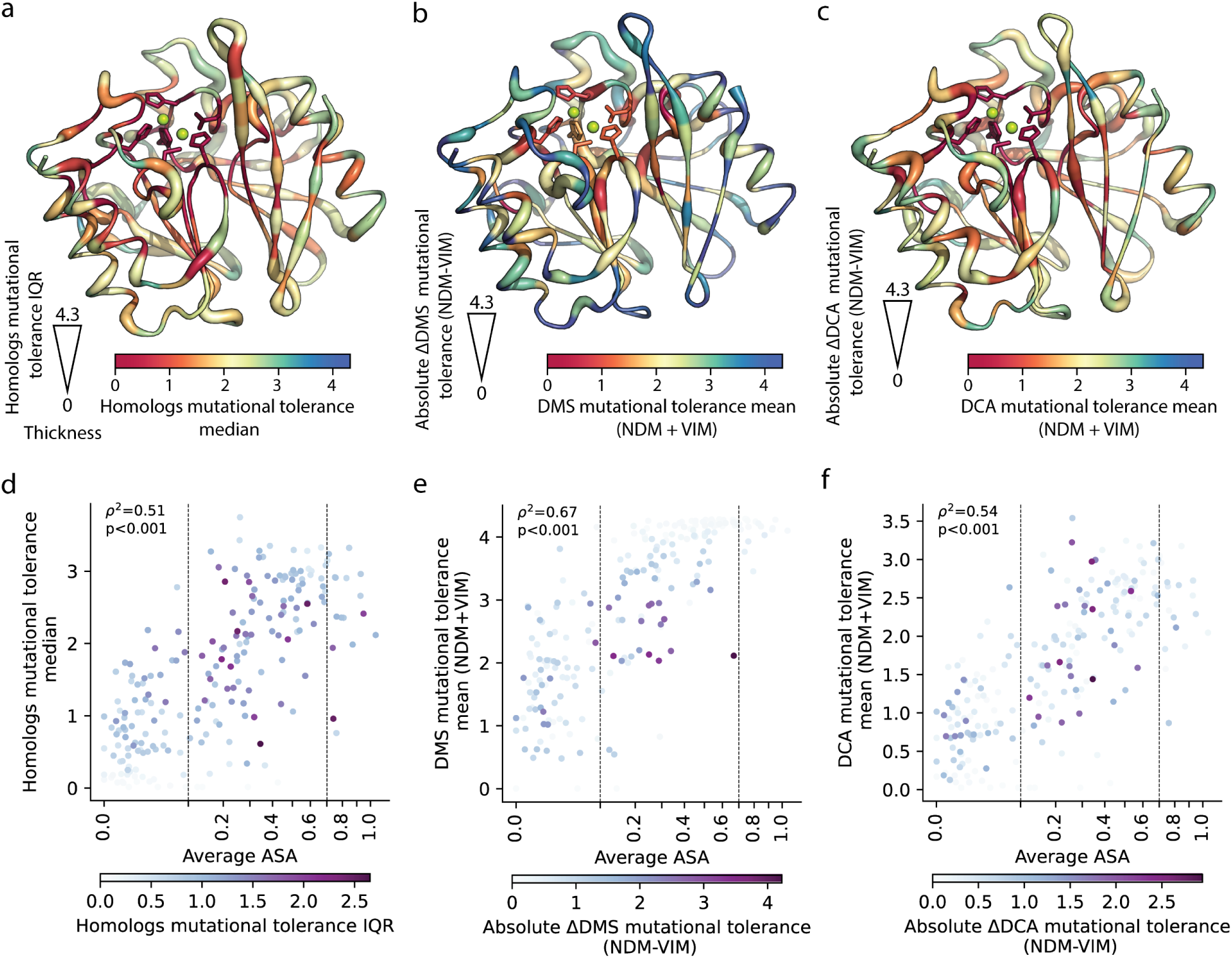
Structural basis of tolerance classifications. (**a**) All homologs mutational tolerance data overlaid on the crystal structure of VIM-2 (5yd7), with the thickness of the backbone representing the IQR in the 100 homologs, and colored by the median. **(b)** DMS mutational tolerance data overlaid on the crystal structure of VIM-2, with the thickness of the backbone representing the absolute difference between NDM-1 and VIM-2, and colored by the average mutational tolerance. **(c)** DCA mutational tolerance of VIM-2 and NDM-1 overlaid on the crystal structure of VIM-2, with the thickness of the backbone representing the absolute difference, and colored by the average. **(d)** Scatter plot of the median mutational tolerance values of 100 homologs versus the average ASA of VIM-2 and NDM-1, with positions colored by the mutational tolerance IQR. **(e)** Scatter plot of the average DMS mutational tolerance of NDM-1 and VIM-2 DCA CDE versus the average ASA, with the positions colored by the difference in mutational tolerance between NDM-1 and VIM-2. **(f)** Same as panel (e) but for the DCA predictions: scatter plot of the average mutational tolerance of VIM-2 and NDM-1 versus their average ASA, colored by the mutational tolerance differences.

We first visualize the mutational tolerance of the 100 homologs superimposed on the structure in terms of the variability (IQR, the thickness of the backbone) and average behavior (median of mutational tolerance, the color scale) (**Fig. 3a**). To compare this global predicted trend with the specific behavior of NDM-1 and VIM-2, we produce equivalent figures using the DMS (**Fig. 3b**) and DCA (**Fig. 3c**) mutational tolerances. We see similar tendencies in the three figures. Regions that are buried in the protein core, including the active-site metal-binding residues, tend to have both low mutational tolerance and spread; likely as a result of being critical to folding or activity and hence mutationally constrained. The most exposed positions generally have high entropy and very low spread according to both the DMS and the DCA, which is a result of both NDM-1 and VIM-2 being completely mutationally tolerant at these positions. Finally, we also observe that some of the more buried residues have higher entropy as computed from the DMS of NDM-1 and VIM-2 than in the DCA model. This observation is consistent with the fact that DCA, which predicts a Gaussian-like distribution of mutational effects^37^, typically tends to underestimate the number of neutral (or almost neutral) mutations that are often one peak in a bimodal distribution^4,6,8^. It can also possibly be due to specific differences in residues and spatial arrangements between the two homologs, as compared to the global distribution of the DCA model.

To quantify the observations about the role of the structural position we use the average accessible surface area (ASA) of NDM-1 and VIM-2. The first pattern that emerges is a significant correlation between average ASA and the site-specific mutability, evident in both experimental data and model predictions (**Fig. 3d-f**). In particular, there is a pronounced correlation between average ASA and DMS derived entropies (Spearman squared = 0.67) (**Fig. 3e**). This observation has been previously reported^4,8,40–42^, and is possibly due to the higher prevalence of structural interactions among internally situated protein residues, thereby amplifying the potential for mutations with adverse effects and vice versa. The correlation is still large (Spearman squared 0.51-0.54) when we compare the ASA to the DCA derived mutability, both at the family level (**Fig. 3d**) and specific to NDM-1 and VIM-2 (**Fig. 3f**). In this case, the capability of DCA to identify these signals is mainly attributable to the conservation patterns embedded within the MSA utilized to train the model.

Furthermore, we used the ASA as a structural variable to distinguish three classes of residues: very buried (ASA<0.1), partially exposed (0.1<ASA<0.7) and very exposed (ASA>0.7). We analyzed in detail how mutational heterogeneity is influenced by the three levels of residue burial. First of all, very buried residues tend to be more mutationally constrained, *i.e.* positions with very low ASA (ASA < 0.1) have low entropy and low spread (blue points in the bottom left of **Fig. 3d-f**). Second, the very exposed positions (ASA >0.7) typically display high mutational entropy and low spread, i.e. they are very mutable in all homologs. These first two observations are consistent with a classical picture of conservation due to structural constraints. Most importantly, we find an intermediate region of partially exposed residues (0.1<ASA<0.7) showing some residues with very high spread in mutational tolerance in DMS and DCA (**Fig. 3d-f**). We also observe that in very buried residues, some positions do exhibit a fairly large mutational variability between the two wildtypes VIM-2 and NDM-1, as highlighted in panels **d** and **e** of **Fig. 3**. Interestingly, this contrasts with the almost total absence of mutational heterogeneity observed between very exposed residues of VIM-2 and NDM-1, especially in the DMS data.

A possible cause of this difference is the numerous intramolecular interactions occurring around buried residues, suggesting that a rich intramolecular network not only reduces the mutability of residues, but it also leads to homolog-specific differences of such constraints. However, it is the intermediately exposed region that exhibits the largest variability in behaviors for all datasets. This ASA range corresponds to positions that have more freedom to mutate than the fully buried positions while still being capable of forming interactions with other residues. The possibility of a mutation is therefore strongly dependent on the sequence context and is, therefore, homolog-specific. The large spread in the mutational patterns of the partially exposed region (**Fig. 3d-f**) is common to all analyses, confirming the dependence between epistatic networks and the structure.

### Epistasis from individual variants

We now turn to the study of intramolecular interactions and use both experimental and model information to characterize the epistatic networks and reveal the interdependencies between residues. For this purpose, we compare the effects of individual variants in the mutational scans of VIM-2 and NDM-1 as fitness differences (δδf = *f*_NDM-1_ – *f*_VIM-2_). A large number of variants can be compared directly for positions in which VIM-2 and NDM-1 have the same WT amino acid, which we refer to as ‘shared’ positions. However, when the WT amino acids are not identical, that is at ‘differing’ positions, a direct comparison is not always possible. Therefore, we devise a systematic classification scheme to analyze mutations in ‘shared’ and ‘differing’ positions between NDM-1 and VIM-2 accordingly (**Fig. 4a**).

**Fig. 4.**
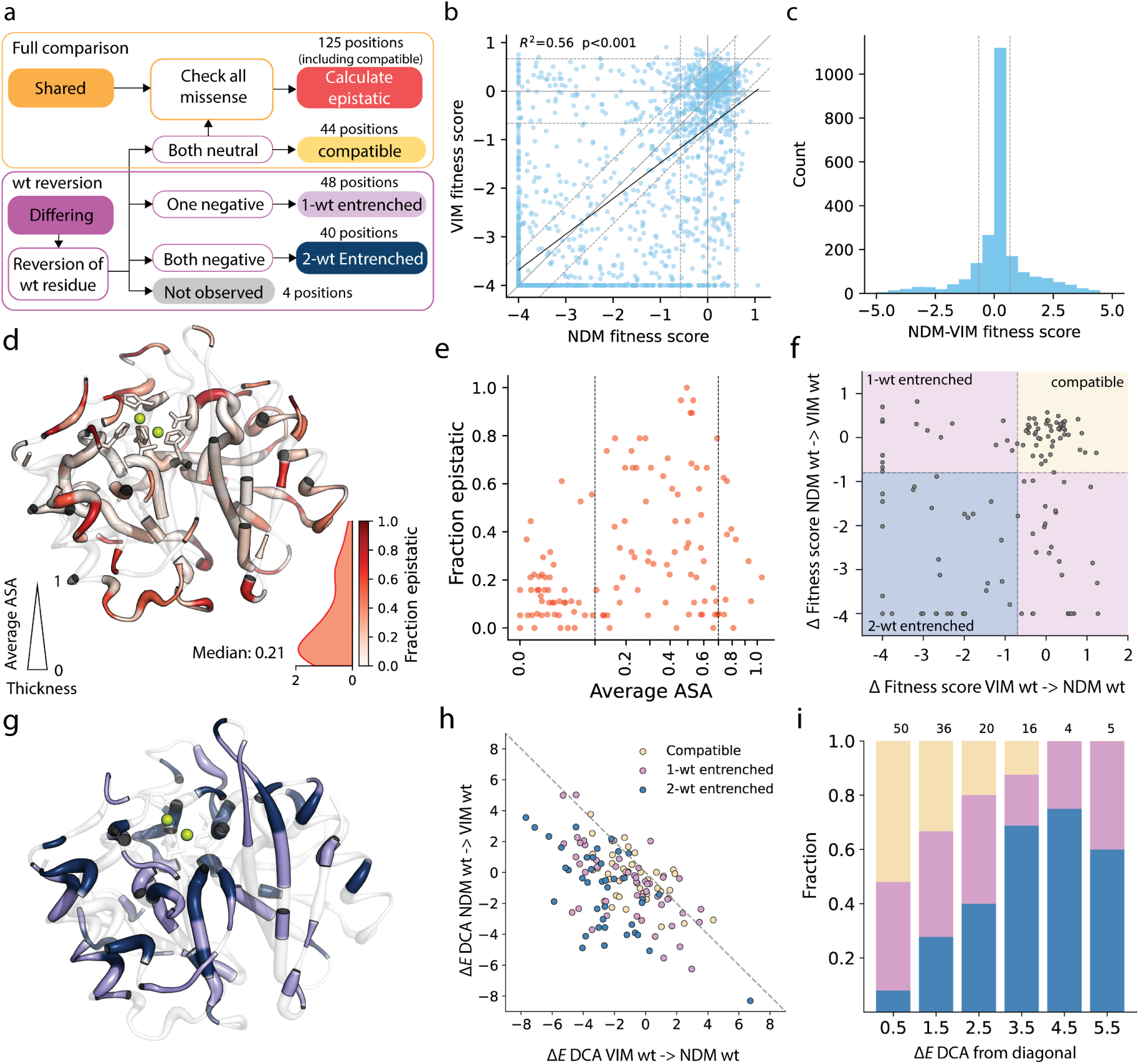
Residue level epistasis between NDM-1 and VIM-2. (**a**) Flowchart of the epistasis classification method. (**b**) Correlation of DMS data between NDM-1 and VIM-2 at shared and compatible positions. The regions between dashed lines in each axis represent the range of neutral fitness effects for each homolog (1.96 x SD of synonymous variants). The diagonal line shares the same neutral range as the y-axis. (**c**) Distribution of fitness effect differences between NDM-1 and VIM-2 at shared and compatible positions. The region between vertical dashed lines represents the range of neutral fitness, equal to 1.96xSD of synonymous variants for NDM. (**d**) Fraction of epistatic mutations overlaid on the VIM-2 structure (shared and compatible positions), and colored by the color scale to the lower right (with distribution). Thickness of the structure corresponds to the average ASA of the NDM-1 and VIM-2 crystal structures. Regions outside the classification are transparent. (**e**) Plot of the fraction of epistatic mutations at each position versus the ASA. Only positions highlighted in panel (**d** are included. (**f**) Scatter plot of fitness effects for mutations of wtVIM-2 towards NDM-1 WT amino acids (x-axis) and wtNDM-1 towards VIM-2 WT amino acids (y-axis) for equivalent positions. The vertical dashed line indicates the left side of the region of neutral effects for VIM-2 based on the synonymous variant distribution, and the horizontal dashed line shows the right side of the region of neutral effects for NDM-1. Quadrants with different behavioral classes are colored as in (**a**. (**g**) Positions that are differing between NDM-1 and VIM-2 that have undergone epistasis analysis of WT reversion mutations. The structure thickness corresponds to average ASA as in panel (**d** and regions outside the classification are transparent. (**h**) Scatter plot of Δ*E* for mutations of VIM-2 amino acids towards NDM-1 amino acids (x-axis) and NDM-1 amino acids towards VIM-2 amino acids (y-axis) for equivalent positions. Points are colored according to their experimental classification as in **Fig 4c**. The dashed line represents the expected behavior without epistasis. (**i**) Bar plot showing the relative fraction of points in each epistatic class at various distances from the diagonal of (**h.** The total number of points in each distance bin is written over the bar.

At ‘differing’ positions, the effect of the starting point of each mutation may impede a straightforward comparison and characterization of epistasis. To accommodate this, for each of these positions we first examine the effect of swapping WT residues between NDM-1 and VIM-2 in both backgrounds. If the swap is neutral in both directions, we define the site as ‘compatible’ (44 positions) and proceed to the comparison of all the missense variants as if the starting points at that position were ‘shared’ (**Fig. 4a**). If the swap is incompatible, that is if one of the mutations is deleterious in either of the two backgrounds (88 positions) we only compare the reversion mutations and consider the position as ‘entrenched’ (**Fig. 4f, Supp.** Fig 3a). We further classify the entrenched positions as ‘1-wt entrenched’ if the swap is incompatible in one background (48 positions) and ‘2-wt entrenched’ if the mutation is incompatible in both directions (40 positions). Because 4 positions lack reversion mutants in our DMS data, we exclude them from the analysis and we label them as ‘not observed’. The great number of entrenched positions points to a pervasive presence of epistasis: for each individual WT ∼50% of WT-swapping mutations (66/132 in NDM-1, 57/132 in VIM-2) lead to a significant loss in fitness, even though they occur naturally in another homolog. Thus, a complex set of compensating epistatic effects must be considered to account for the collective presence of those mutations.

We then analyze ‘shared’ positions, that is positions with the same WT amino acid. Together with the ‘compatible’ positions described earlier, they add up to a total of 125 residues which we collectively call ‘comparable’. The analysis of the DMS data for these positions allows us to compare equivalent mutations across the NDM-1 and VIM-2 backgrounds and, therefore, directly quantify context dependence. In this case, we can compare the fitness of all mutations A->B, where A is the common WT amino acid and B represents one of the 19 possible mutations (18 for ‘compatible’ residues). Mutational effects are strikingly different between homologs, with an overall Pearson correlation *R*^2^ of 0.51 (**Fig. 4b**). For each mutation, we consider it epistatic if the fitness difference δδf is greater than the range of neutral effects (standard deviation [SD] of the synonymous variants of NDM-1). A large proportion of variants (∼55%) shows a statistically significant difference of mutational effects between homologs; differences are not strongly biased in sign or effect size toward a single homolog (**Fig. 4c**). To evaluate the degree of epistasis, for each position we compute the fraction of epistatic mutations. We observe a widespread presence of epistasis (**Fig. 4d**) across the entire structure. Quantified as a distribution, we find that the median fraction of epistatic mutations at each position is 0.21, meaning that half of the positions have significant epistasis in at least ∼4 mutations (0.21 x 19 missense aa). The fraction of epistasis at each position can be used to compare site-specific epistasis of VIM-2 and NDM-1 with the spread in mutational tolerance across all 100 homologs and ASA (**Fig. 4e**). Strongly epistatic positions are enriched in partially exposed positions, as previously noted (**Fig. 3d-f**).

The subset of positions classified as entrenched is especially suited to study the influence of epistatic constraints in evolution. They cannot be easily linked to site-specific features or residue burial, as they are scattered across the whole protein structure (**Fig. 4g**); though we do find a subtle bias for 1-wt entrenched positions to be less buried than 2-wt entrenched positions (**Supp.** Fig 3b). Using DCA, we infer the deleterious effects to be caused by the sum of many distributed negative interactions with the surrounding amino acids. This picture points to a model of protein evolution where the effect of mutations changes gradually due to the incremental accumulation of small-magnitude interactions between residues^43^.

We can exploit a DCA-based analysis at the level of individual mutations to explore the role and meaning of entrenched positions in VIM-2 and NDM-1. We produce a scatterplot analogous to that of **Fig. 4f**, but using Δ*E* instead of fitness (**Fig. 4h**). Mutations mainly concentrate around the neutral center, or populate the lower half-plane defined by the diagonal. As expected, we see almost no points towards the upper half-plane, where the swap of both WTs would result in higher fitness. In the absence of epistasis, the effect of the mutations A->B and B->A in different backgrounds should have opposite signs, exhibiting a perfect anti-correlation of the reversion effects, indicated by the diagonal in **Fig. 4h**. In contrast, the further away the points are from the diagonal, the greater the difference in mutational effect in the two sequence contexts. We verify this idea by using the distance from the diagonal as an “entrenchment metric”. We compute the fraction of mutations from each entrenchment class at different distances from the diagonal (**Fig. 4i**). We see that regions far from the diagonal are enriched for 2-wt entrenched mutations, those near the diagonal are mostly compatible, while intermediate regions have mutations with mixed classifications.

The biggest qualitative difference between the mutation reversion plots in **Fig. 4f** and **Fig. 4h** is that DCA predicts many WT-swapping mutations to be beneficial, in clear contrast with the DMS data. We believe this to be a limit of the antibiotic selection assay, where a threshold like relationship between resistance and fitness for a given antibiotic concentration means variants with greater than needed resistance do not necessarily have higher fitness^6^, *i.e.*, we have limited detection for gain of function. As a consequence, epistatic and non-epistatic mutations could be confounded for each other, as illustrated in **Supp.** Fig. 3c. The 2-wt entrenched mutations are guaranteed by the experiment to be epistatic. However, for 1-wt entrenched mutations, if a reversion negative in one background is neutral in the other direction, it could be truly neutral (epistatic), or a beneficial effect that could not be detected (non-epistatic). There are a few reasons that suggest that indeed most of those mutations are truly neutral in one background and deleterious in the other.

First, the interpretation is supported by the distribution of effects observed in our previous analysis of VIM-2 variant EC_50_, which unlike fitness, is not limited by the threshold effect. The rarity of gain-of-function mutations (constituting only 1-2% of all occurrences) makes it improbable that there would be sufficient data points to create the anti-correlation expected in the absence of epistasis. Moreover, DCA supports this interpretation as well: as we have shown in **Fig. 4i** many 1-wt entrenched positions are statistically different from compatible ones, to the extent that some reach far away from the diagonal, just like 2-wt entrenched mutations. We argue that, once again, the model and the experiment complement each other: the model suggests that epistatic interactions identified in the experiment are sparse and pervasive and supports the interpretation of 1-wt entrenched mutations as being mainly neutral in one of the two directions. Moreover, DCA proves to be quite accurate in predicting epistasis: all of the most distant points from the diagonal are either 1-or 2-wt entrenched according to the experiments.

### Probing specific epistatic interactions

A subset of our data also gives us the opportunity to dissect specific epistatic interactions in the homologs. Positions that have entrenched WT provide a potential signal for residues that are functionally important and may participate in specific interactions. We sought to find surrounding interactions within the proteins by identifying pairs of positions that are in close proximity, and where the pair of positions have entrenched WT residues (**Supp.** Fig. 4a). One example of shape complementarity can be found between NDM-1 positions 125 and 249 (VIM-2 positions 119 and 239) directly underneath the active site Zn^2+^ ions, where both are entrenched positions within contact distance to each other (**Fig. 5a**). In VIM-2, the Arg at position 125 is paired with Gly at position 249. In contrast, NDM bears a somewhat smaller, but still positively charged Lys at position 125, which is now paired by a larger Ser at position 249. To test if these positions possess significant interactions, we generated the single and double mutants of each position pair in the NDM-1 background by mutating to the VIM-2 WT at those positions, then measuring their phenotype (*EC*_50_ against AMP) (**Fig. 4c**). As the effect of VIM-2’s WT is deleterious in NDM-1 as single mutants, we expect that mutating all interacting positions may lead to compensation, generating a less negative effect. We also extend this experiment to pairings in the L3 active site loop (NDM-1 positions 67+68), L10 active site loop (218+266, 211+229) and some buried positions beneath the L10 loop (197+204, and a triplet of 204+246+259) (**Fig. 5c**); the triplet was tested as all pairs of doubles and the full triplet. Combined, our selection of positions covers a variety of positions, including those with different biochemical properties (size, polarity, charge), different solvent accessibilities and different spread in mutational behaviors across the whole family (**Fig. 5c, d**).

**Fig. 5.**
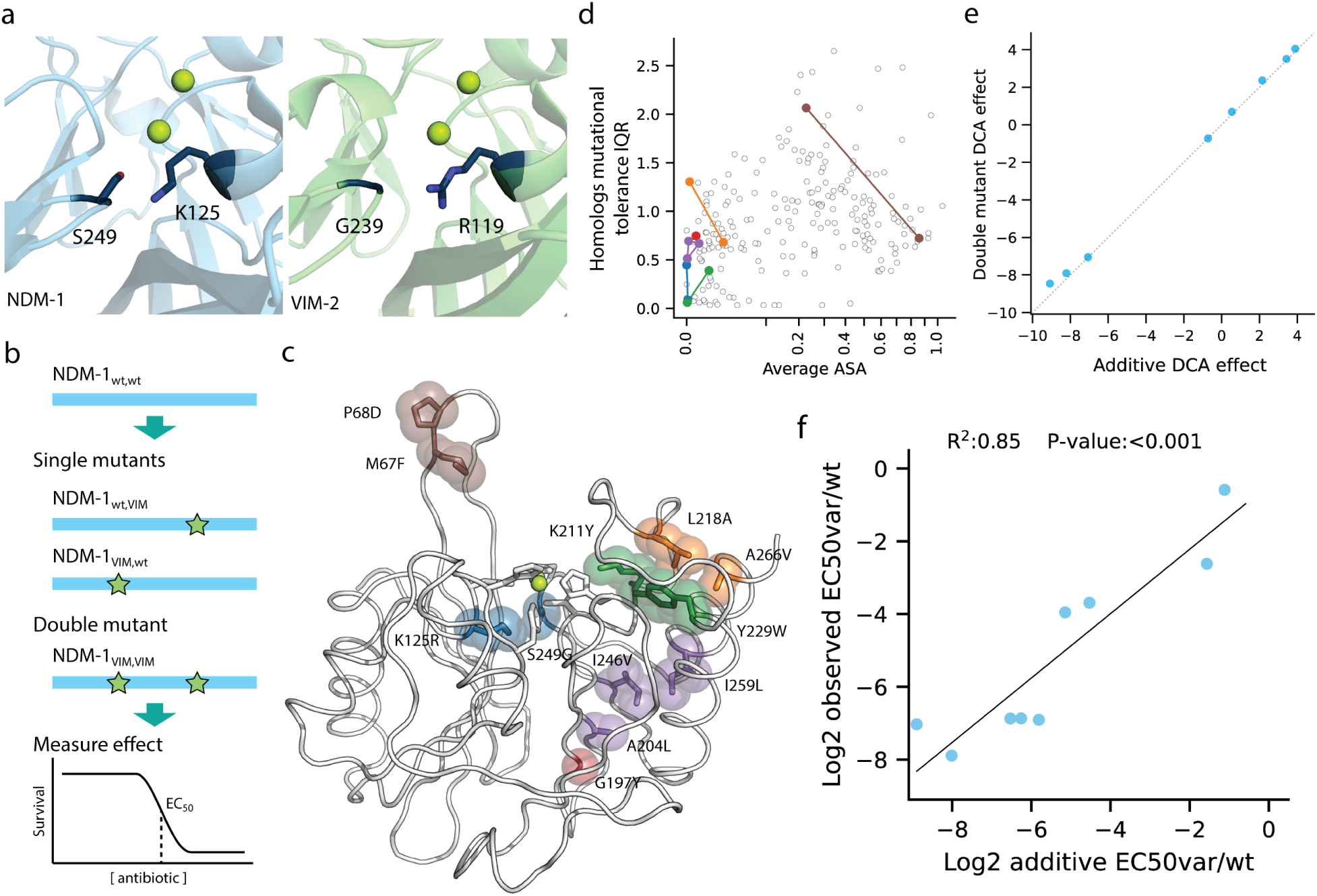
Testing interactions of entrenched positions in NDM-1. (**a**) Example of potentially interacting entrenched WT positions in the crystal structures of NDM-1 and VIM-2. (**b**) Experimental scheme for testing single or combined mutational effects in the NDM-1 background. (**c**) Entrenched WT positions that were chosen for testing of epistatic interactions. Positions with the same color are mutated together to test for compensation of entrenchment; A204L overlaps 2 sets and is also tested with G192Y (red). (**d**) Plot of IQR in CDE across 100 homologs and the average ASA of NDM-1 and VIM-2 structures, with the tested positions highlighted. Tested combinations are shown as lines. (**e**) Scatterplot of DCA energy change of all selected double (1 triple) mutants, with the expected additive single mutant effects in the x-axis, and the observed double mutant effects in the y-axis. (**f**) Scatterplot of all tested double (1 triple) mutants, with the expected additive single mutant effects in the x-axis, and the observed double mutant effects in the y-axis. Effects are calculated as fold change relative to wtNDM-1.

The effects of the single mutants validate the deleterious nature of the observed fitness scores, and we observe a sigmoidal relationship between a mutant’s *EC*_50_ and the fitness score, which is consistent with previous observations^6^ (**Supp.** Fig. 4b). This validates the entrenchment observed in DMS, as all selected mutations that are WT residues in VIM-2 are deleterious in the NDM-1 background. It is also notable that our selected mutations evenly span a wide range of deleterious effects. When we tested the mutants in combination, however, we did not observe significant compensation in any of the mutant combinations (**Fig. 5f**). In fact, the log-additive effects of the single mutants (null model for no epistasis) show a distinctly linear correlation with the observed double mutant effects, with an *R*^2^ of 0.85. It appears that entrenched positions cannot be easily swapped simply by mutating other nearby entrenched residues. This scenario is confirmed by DCA where, as previously discussed, the epistatic effects can only arise as a cumulation of small contributions including a multitude of epistatically coupled positions. Previous experiments have also shown that direct compensation by nearby residues is expected to be rare^13^, and compensation often requires more than two mutational steps^44^. The double mutation effects are therefore basically additive in DCA (**Fig 5e**) as in the experiment (**Fig 5f)**. It is also not the case that the positions are globally independent in their effect, as these same mutations are fixated in VIM-2 and the deleterious effects have been compensated for during evolution. Thus, these experiments suggest that the intramolecular network of residue interactions is much more complex, and even entrenched WT residues are not themselves sufficient to dictate the epistatic networks.

## Discussion

By combining the computational approach of DCA and the experimental approach of DMS, we have gained a global picture of the epistatic tendencies between the homologs within a protein family. In the B1 β-lactamase family, we find a prevalence of heterogeneity in mutational effects at both the level of overall mutational tolerance at each position, as well as epistasis of the same mutation across different backgrounds. At a global level across all homologs, over half of the positions can exhibit a >6 fold difference in mutational tolerance depending on the sequence background. When examined specifically in the NDM-1 and VIM-2 contexts, half of positions with shared WT residues have >4 mutants that are epistatic. In particular, the complementarity of the two approaches has enabled better understanding than either approach alone. The global level view provided by DCA for 100 homologs reveals the full spread in mutational behaviors, which would have been obscured for positions that behave similarly for the 2 specific homologs tested. In turn, while both approaches generally agree on the mutational behaviors of each position, the experimental results can highlight peculiarities that the statistical approach may not identify from just the evolved sequence data.

To expand upon previous DMS studies involving multiple homologs, which are generally focused on higher-level statistical analyses^26–29,31,43^, we also performed deeper examination of the mechanisms behind the observed incompatibilities. The ASA of a protein position shows trends with various facets of epistasis, as global spread is more constrained in behavior at extremely low or high ASA, while epistatic behavior between two specific sequences seems most prevalent at intermediate ASA. These observations are likely underpinned by differential intramolecular networks in different homologs, as a result of gradually co-evolved epistatic networks over the course of evolution. However, directly replacing just two or three residues is not enough to compensate for evolutionarily entrenched residues, suggesting that a much more complex network of interactions is at play^45^, as predicted by the DCA model.

We suspect that many of the observed behaviors are likely not limited to NDM-1 and VIM-2, or the B1 family. The trends with regards to structure in both DCA and DMS, which are typical of those observed in other systems^4,46–49^, suggest that the behaviors we observe can be explained through general mechanisms such as secondary structure formation and structural packing. However, more evidence from other systems would be required to distinguish the global trends from system-specific trends. Overall, we provide a global view of epistatic networks at the protein family level that complements more detailed examinations of epistasis in specific residues^29,50^.

## Methods

### Sequence collection for the B1 MBL family

In an effort to comprehensively isolate all B1 MBLs, we used a broad sweep approach through a sequence similarity network (SSN) using data from genomic (UniProt) and metagenomic (JGI) databases. Initially, an SSN was constructed using the EFI-EST tool using the InterPro ID IPR001279 (MBL domain superfamily) as the input. The network was analyzed at different alignment score cut-offs to find a cut-off where all clinically isolated B1 enzymes fell within the same isolated cluster (∼1,800 sequences at EFI alignment score cut-off of 25). The sequences that were shorter than 220aa were removed, and the remainder (∼1,300) were clustered by cd-hit to 60% identity to reduce redundancy (187 sequences final) then used to generate an HMM, *i.e.*, sequence profile, of the B1 family. The HMM was used to search for B1 sequences in the JGI database resulting in ∼2.5Mil sequences from JGI, with 1,859,503 non-redundant sequences determined by cd-hit clustering at 100%.

To construct an initial SSN, all sequences of the MBL domain superfamily were downloaded from UniProt using the InterPro IPR001279 definition (434,623 sequences, accessed 2019). The UniProt and JGI data were combined, sequences with ambiguous characters were removed, and the length was restricted to between 100-1500aa. Cd-hit was used to successively cluster the sequences to 100%, 90%, 70% and finally 50% identity to reduce redundancy, resulting in 86,947 representative sequences for the entire MBL superfamily. A SSN was generated by performing ‘all by all’ BLASTP with an e-value cutoff of 1e-5 and 1000 maximum hits per sequence, giving a raw network of ∼90Mil pairwise bitscore values (edges) between all sequences (nodes). The network was processed using an inhouse SSN analysis pipeline (MetaSSN: https://github.com/johnchen93/MetaSSN) to identify the lowest bitscore cutoff at which all clinically isolated B1 sequences break off into an isolated cluster (10,252 sequences). Finally, to identify sequences that are most likely to be active B1 sequences, the dataset was filtered by length to between 200-350aa (6,828) and used to build a multiple sequence alignment (MSA) using Clustal Omega on default settings. The MSA was manually curated to exclude any mis-aligned sequences, resulting in 6308 curated sequences. Finally, to ensure the sequences were likely to have B1-like function, only sequences aligned with the conserved B1 active-site metal binding residues (H116/H118/H196 and D120/C221/H263, BBL numbering) were kept, resulting in a final dataset of 5035 B1 family sequences, with a roughly 50/50 split between sequences from UniProt and JGI.

### Direct coupling analysis of the B1 family

To generate an alignment of conserved structure, crystal structures of 14 homologs were aligned using mTM-align (https://yanglab.nankai.edu.cn/mTM-align/)^51^. The homologs (PDB IDs) are: NDM-1 (3spu), VIM-2 (5yd7), DIM-1 (4wd6), ECV-1 (6t5k), FIM-1 (6v3q), GIM-1 (2ynt), IMP-1(4uam), IND-7 (3l6n), MYO-1 (6t5l), TMB-1 (5mmd), VMB-1 (6jv4), bc-II (1bc2), blaB (1m2x) and cfiA (1znb). An HMM sequence profile was trained on this curated dataset by using HMMER via the *hmmbuild* command. We used a –*symfrac* value of 0.3 to control the maximum number of gaps in each alignment column. The 5035 B1 sequences were then aligned to this profile via the *hmmsearch* command to produce the MSA for training the DCA model. After converting the alignment to FASTA format, all sequences exhibiting more than 10% gaps were removed from the alignment. Additionally, flank columns exhibiting more than 75% gaps were eliminated. The resulting alignment contained 222 sites. A final refinement was achieved after removing all sequences that exhibited more than 80% sequence identity to NDM-1 or VIM-2, thereby ensuring the alignment was not biased toward the sequences used for further analysis. The resulting MSA had 3655 sequences. We then inferred on this alignment a DCA model by using standard settings of the adabmDCA package corresponding to Persistent Contrastive Divergence using 40 MC sweeps at each iteration (PCD-40)^17^.

### Library generation and deep mutational scanning of NDM-1

The procedure for library generation and DMS on NDM-1 were conducted in an identical manner to VIM-2. The wtNDM-1 sequence is encoded on an inhouse pIDR5.1 plasmid, expressed under a constitutive AmpR promoter. To generate all single amino acid mutants, we used a PCR based method (restriction free cloning^52^) to introduce a degenerate ‘NNN’ sequence at each codon in the coding sequence in separate reactions. After each single position was mutated, the library was combined into 7 groups of 39 consecutive positions each, forming 117nt long mutated regions that can be fully sequenced by paired end Illumina NextSeq reads.

The NDM libraries are then transformed into *Escherichia coli* (*E. cloni* 10G, Lucigen) and stored as glycerol stocks, with the number of colony forming units after transformation measured to be >= 100,000 to ensure complete coverage of each group. To perform selection experiments, we inoculate the glycerol stocks into fresh LB (Fisher) and grow the cultures shaking overnight at 30°C for 16hrs. The cultures are diluted into fresh LB to an inoculum of 1.5×10^6^ cells/mL (targeting a 1:1000 dilution of a culture with OD_600_ of 1.5) and grown shaking at 37°C for 2hrs. Selection pressure is introduced by mixing 960uL of cell culture with 40uL of a 25x stock of each antibiotic suspended in LB, for a final culture volume of 1mL. We tested selection at 32, 128 and 256ug/mL AMP, 2, 16, 32ug/mL CTX, and 0.063, 0.25 and 0.5ug/mL MEM. We also grew a sample of the library without selection. The culture is grown under selection while shaking at 37°C for 6hrs, then removed from selection via centrifugation and resuspension in 1mL of fresh LB, repeated 3 times. The post selection culture is grown, shaking overnight at 30°C for 16hrs and the plasmid DNA is purified using a QiaPrep 96 column DNA purification kit (Qiagen). This procedure was conducted twice on different days to produce two separate replicates.

To deep sequence the selected library, we used primers targeting unmutated regions that directly flank the mutated region of each of the 7 library groups to amplify the DNA and to attach Nextera adaptors to the amplicons. The amplicons undergo a second PCR to attach the Illumina sequencing indices and flowcell binding sites. All tested samples (all groups, conditions, replicates) were sequenced in the same Illumina NextSeq 550 run with fully overlapping paired end reads. We also included control samples of amplicons extracted from just wtNDM-1 using the primers for each group in the NextSeq run. After deep sequencing, the forward and reverse reads are merged together using our previously published pipeline (https://github.com/johnchen93/DMS-FastQ-processing), and we discard reads with greater than 20 mismatches between forward and reverse reads or with a posterior Q score of less than 10. We use the wtNDM-1 samples as an estimate for error rates arising from the deep sequencing process, and we filter the non-selected libraries to remove variants with frequencies less than 2x of the expected frequency from sequencing noise alone, or variants with less than 5 reads. We then calculate the fitness scores for each variant in all conditions according to eq (1). All variants that pass filtering in the non-selected condition are considered to truly exist in the library, and if the same variants are not observed in conditions undergoing selection they are assumed to have been depleted by selection and are given a dummy count of 1 to simulate the lowest possible frequency. Raw sequencing data is available on the NCBI Sequence Read Archive under BioProject PRJNA974578.

### Calculating mutational tolerance

To calculate DCA mutational tolerance, we calculate the probability of each mutant at a given position relative to the WT (proportional to exp(-Δ*E*)).

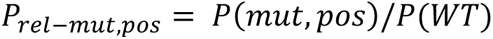

We then calculate the mutational tolerance using the probabilities of all mutants at that position through the Shannon Entropy.

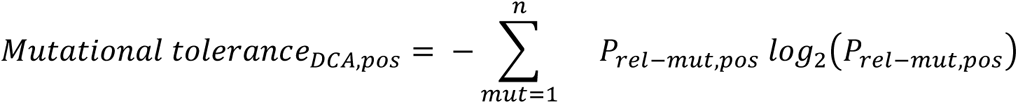

For DMS mutational tolerance, we take an analogous approach to Firnberg *et al.*^4^. First, we normalize our fitness scores from a range of 0 to 1 relative to WT, where beneficial mutations are >1, and deleterious mutations range from 0-1. Since fitness scores of –4 or lower correspond to missense variants, we use –4 as the minimum range of fitness scores. We add a small padding factor of 10^5^ to avoid complete zeroes.

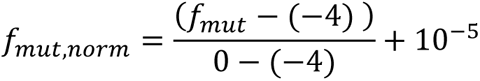

We use the proportion of normalized fitness relative to the sum of all normalized fitness scores as a measure of probability. If the position does not have synonymous mutants, WT is inserted with a normalized fitness of 1.

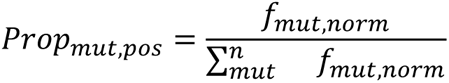

DMS mutational tolerance is then first calculated as the Shannon Entropy.

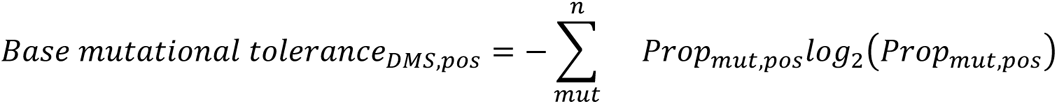

We also scale the base mutational tolerance so that the effective number of amino acids is a maximum of 20, to account for positions which do not observe all 20 amino acids. The result is then the final DMS mutational tolerance.

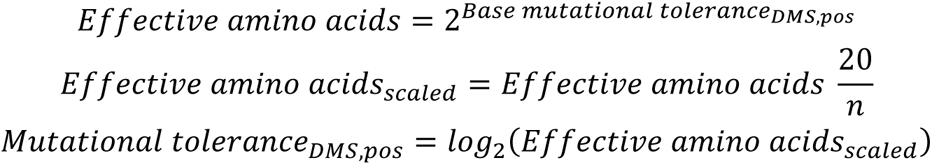

### Generation and dose-response assay of NDM single and combined mutants

The selected positions with entrenched WT behaviors were mutated in the NDM-1 background from the NDM-1 wt residue to the VIM-2 wt residue using Golden Gate cloning as single mutants (NDM-1 positions M67F, P68D, K125R, G197Y, A204L, K211Y, L218A, Y229W, I246V, I259L, A266V). Then, combinations of positions were generated by a second (or third for the triplet) round of mutations. Dose response curves were carried out to obtain the half-maximal effective concentration (*EC*_50_) under ampicillin (AMP) selection. For testing, mutants and wtNDM-1 were transformed into *E. coli* and grown overnight for 16 hrs at 30°C, then diluted to a target inoculum of 1.2Mil cells / mL (OD_600_ of 0.0015) the next day. The diluted culture is grown for 1.5hrs at 37°C, then 180uL of culture is mixed with 20uL of 10x AMP stock, with final ampicillin concentrations from 1-1024 ug/mL. Growth under AMP selection is done for 6 hrs at 37°C, and the OD_600_ of each culture is measured after selection. For each mutant or wtNDM-1, the OD_600_ across all selected AMP concentrations is plotted as a dose-response curve and fitted using a sigmoidal equation to obtain the *EC*_50_.

## Supplementary Data

**Supplementary figure 1.**
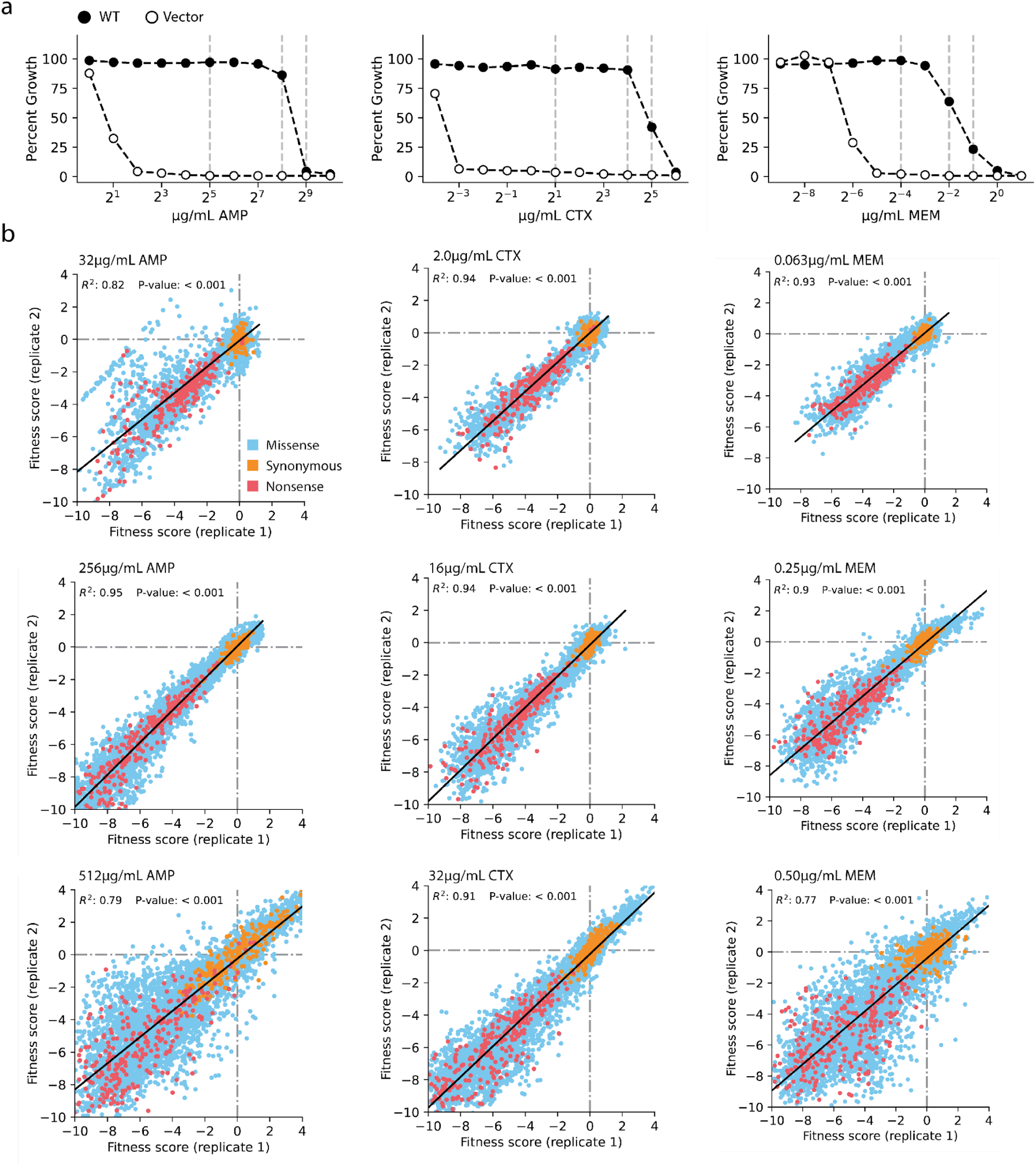
Selection range and replicate correlation of NDM-1 DMS experiments. (**a**) Growth curves for wtNDM-1 and empty vector under selection with 3 different antibiotics. Vertical dashed lines indicate conditions explored in DMS experiments. (**b**) Replicate correlation for NDM-1 DMS data, conducted in the 3 antibiotics at 3 concentrations each.

**Supplementary figure 2.**
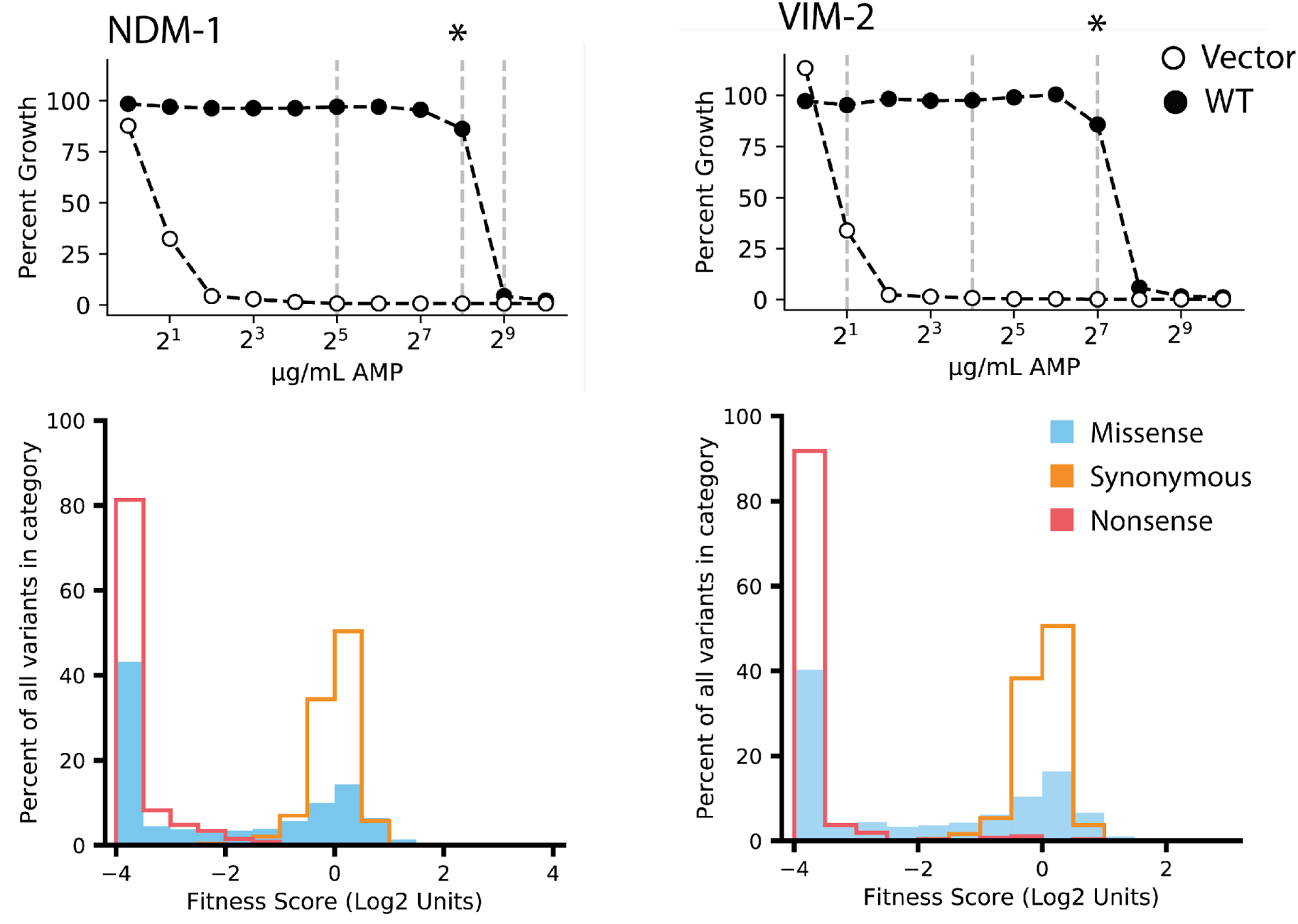
Comparison of selection pressures between NDM-1 and VIM-2 dataset. In the top panels, the dose-response curve shows the survival of E. coli expressing each WT or an empty vector across increasing selection pressures of AMP. The stars indicate the selection conditions of the datasets being compared directly in this study. The bottom panels show the distribution of fitness effects of all mutants of NDM-1 or VIM-2, split into categories of the type of mutation.

**Supplementary figure 3.**
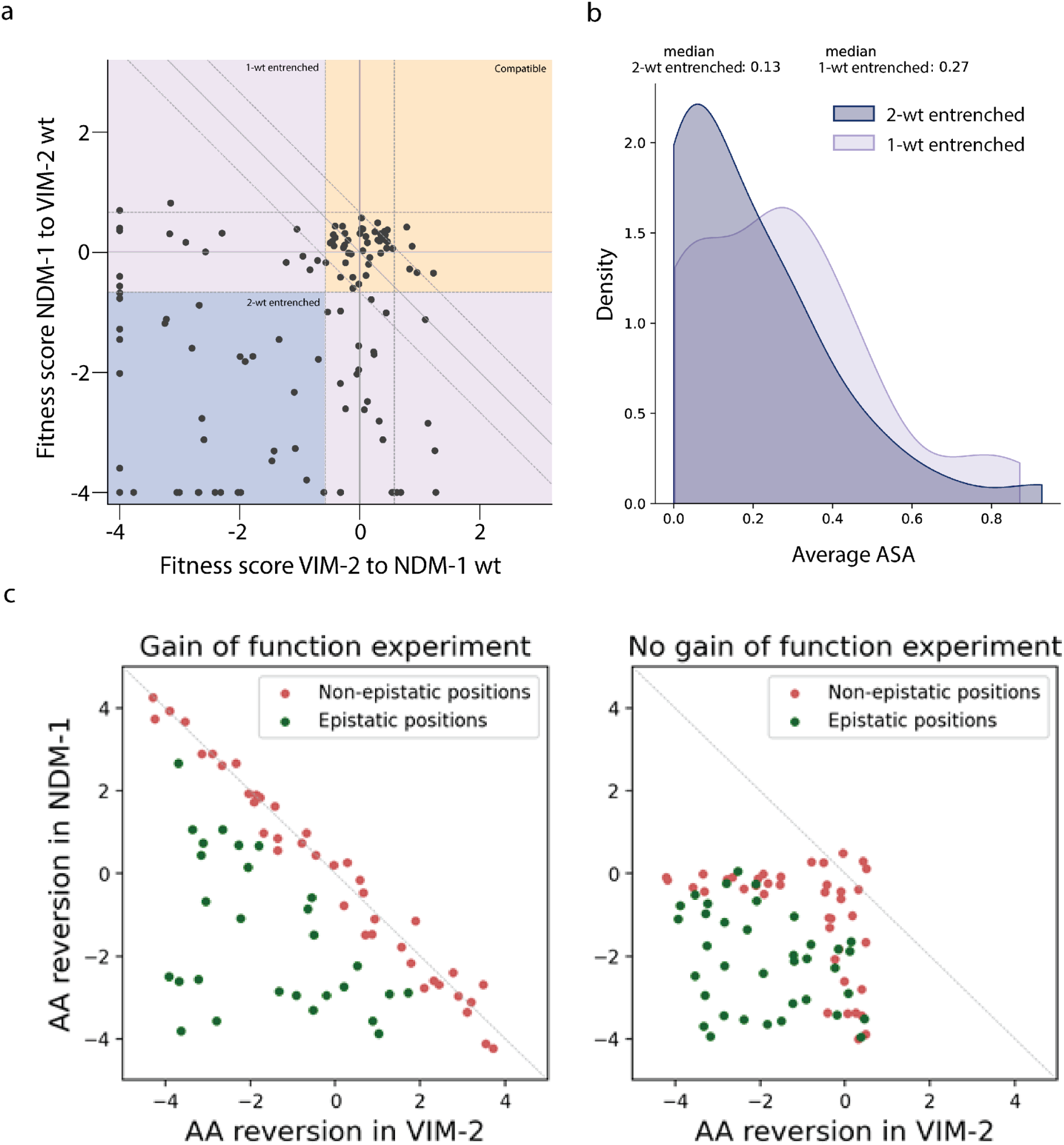
Epistasis of entrenched residues. (**a**) Scatter plot of fitness effects when wtVIM-2 is mutated to the NDM-1 wt residue at a position (x-axis), or wtNDM-1 is mutated to the VIM-2 wt residue (y-axis). The dashed lines indicate the region of neutral effects based on the synonymous variant distribution (1.96 x sd), for NDM-1 (horizontal, diagonal) or VIM-2 (vertical). The diagonal line indicates the expected behavior with no epistasis and no fitness plateau, *i.e.*, each mutation has the same magnitude but opposite sign when reverted. Quadrants with different behavioral classes are colored. **(b)** Distribution of average ASA for positions that are 1-wt or 2-wt entrenched. **(c)** Hypothetical distribution of reversion mutations in an experiment capable of detecting gain of functions (left) or incapable of detecting gain of functions (right).

**Supplementary figure 4.**
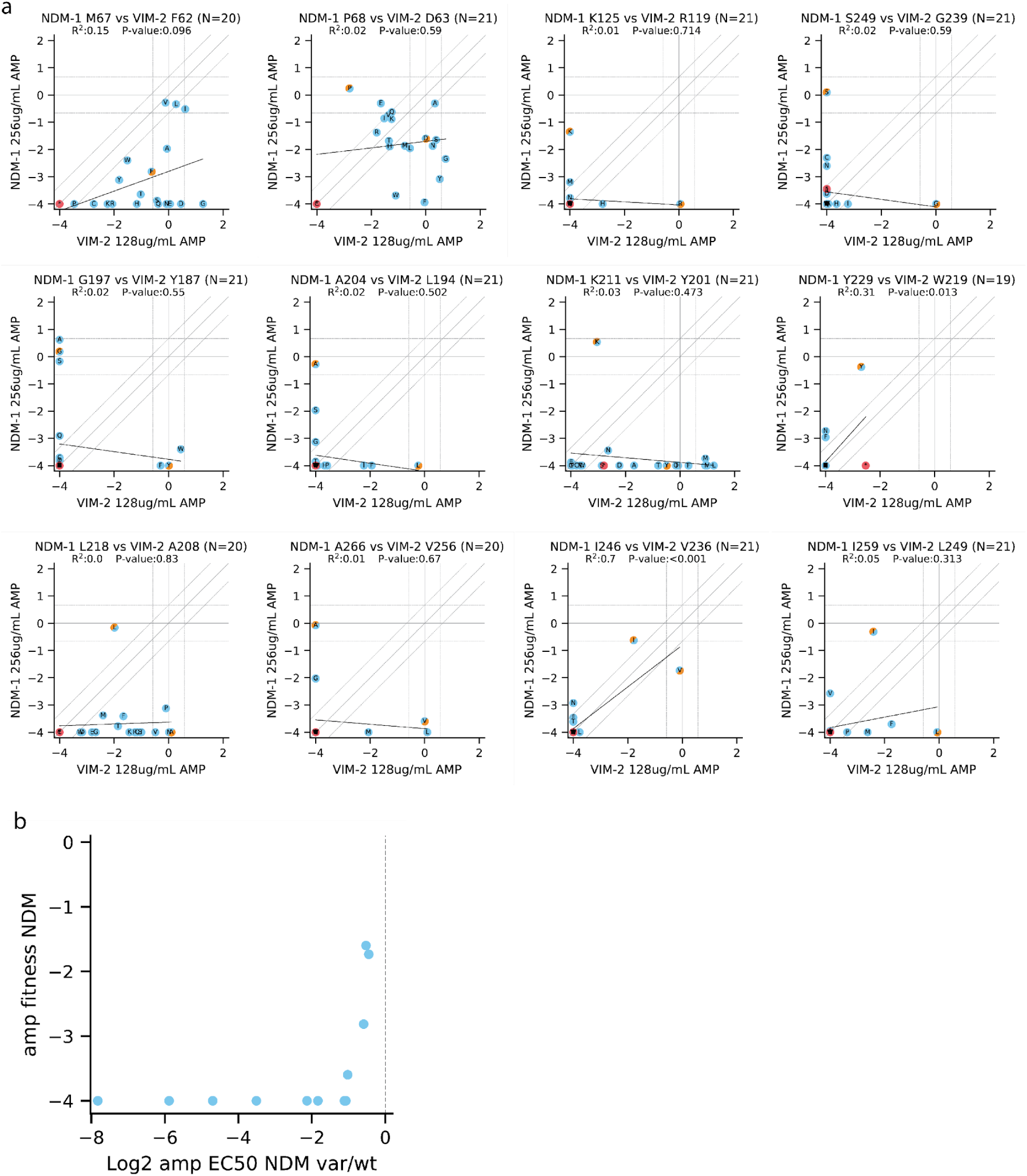
Fitness scores at positions in the NDM-1 background selected for mutation to the VIM-2 wt. (**a**) Scatter plots show the fitness scores of mutants between NDM-1 and VIM-2 at aligned positions (labeled at top of each figure). Points with orange color in one half indicate the synonymous mutant for the background (e.g., orange in lower right side indicates a synonymous variant of VIM-2); some synonymous variants are not plotted (M, W have only one codon), but the mutation of the other background to the wt are still in the dataset. Red points are nonsense variants. Blue points indicate all other missense variants. Vertical and horizontal dashed lines indicate the regions of neutral effects based on the synonymous variant distributions of each homolog. The diagonal dashed lines have the same region of neutral effects as the y-axis. **(b)** EC50 vs DMS fitness of the selected entrenched double and triple mutants. EC50 of each variant is expressed as the fold change from wtNDM-1.

